## Supplementary figures and images for "Rest-task Modulation of fMRI-derived Global Signal Topography is Mediated by Transient Co-activation Patterns"

### S1_Fig.tif

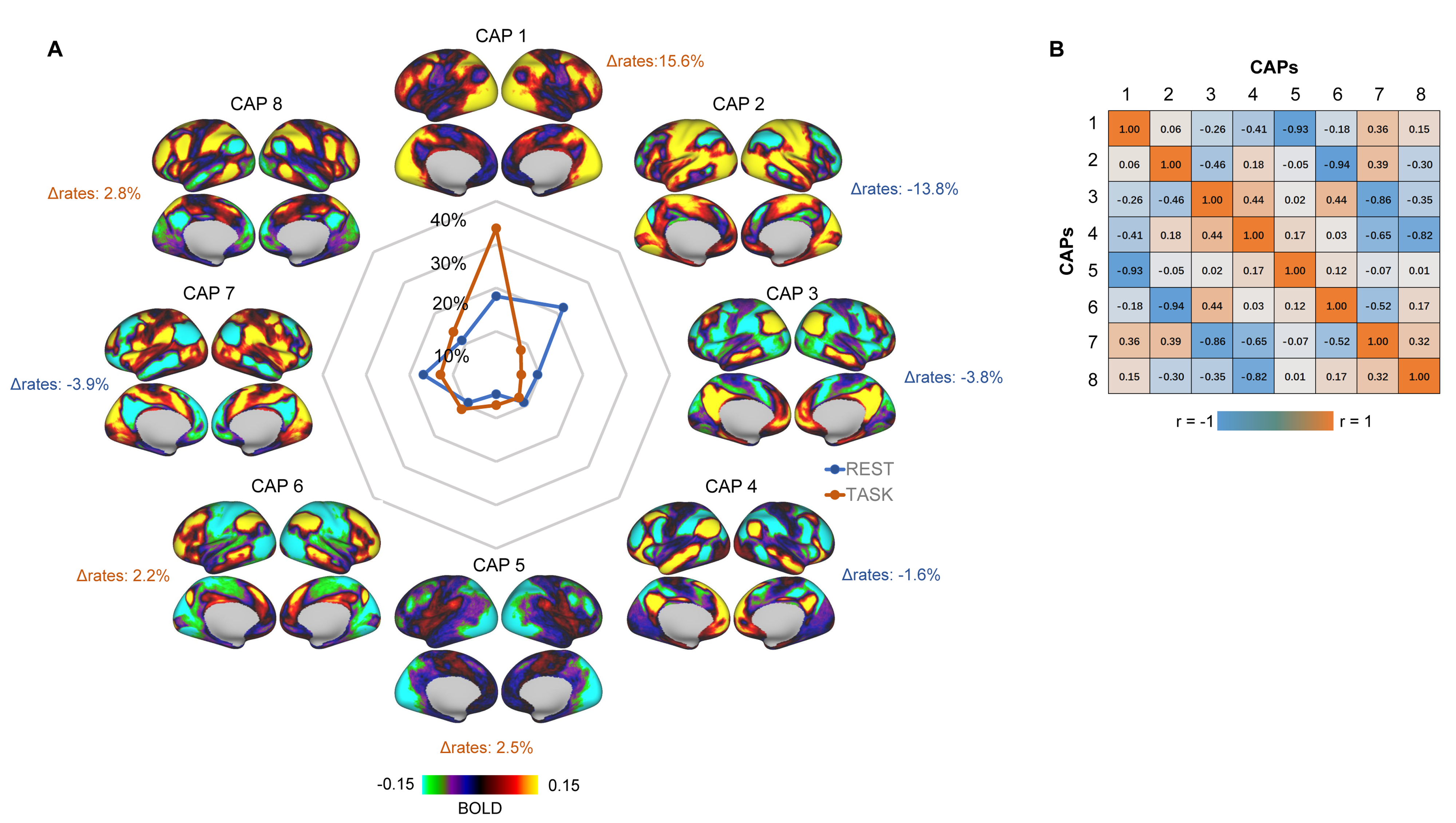

### S2_Fig.tif

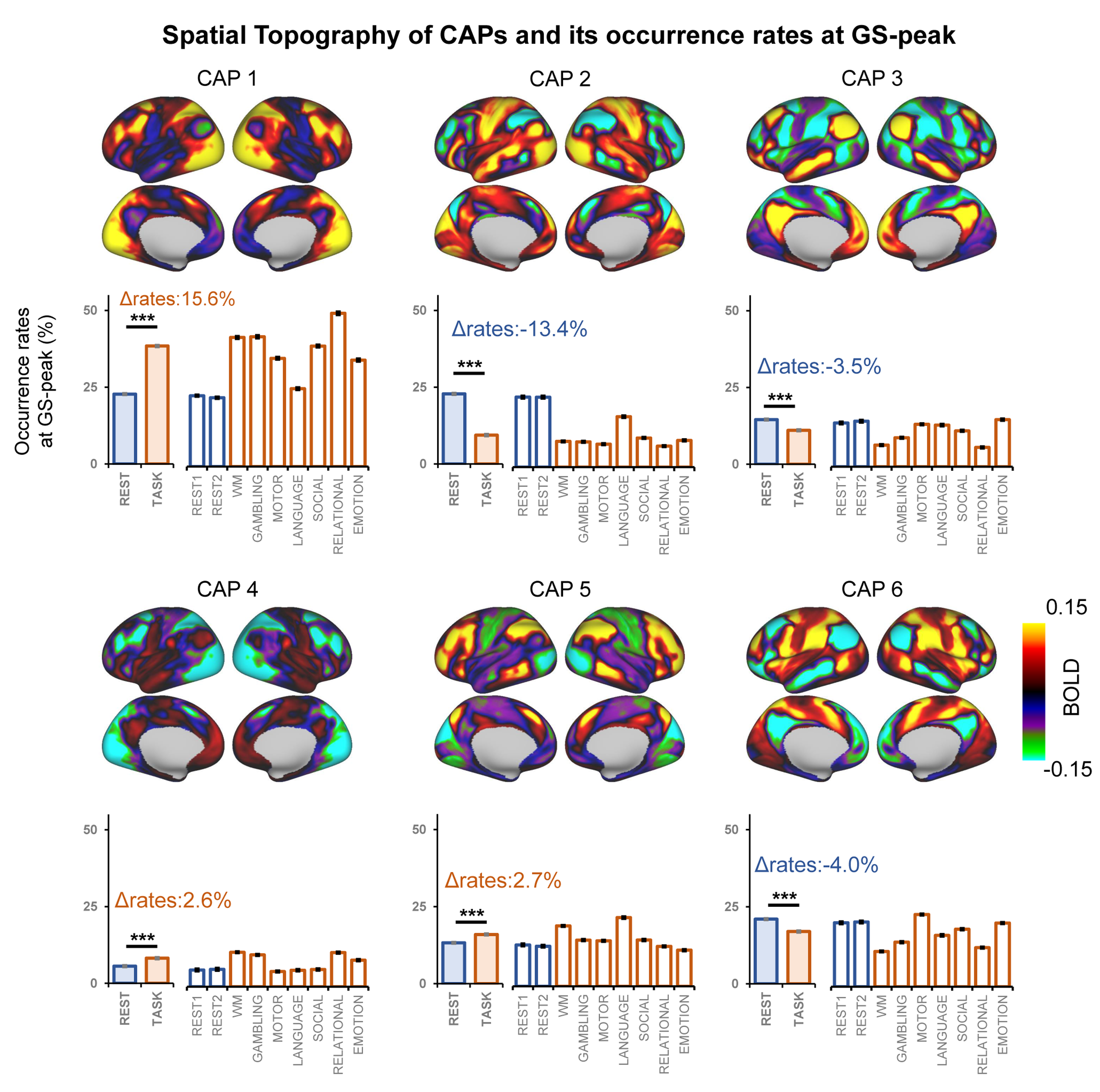
